# On-demand insulin manufacturing using cell-free systems with an “on-column” conversion approach

**DOI:** 10.1101/2025.04.08.633785

**Authors:** Shayan G. Borhani, Max Z. Levine, Chandrasekhar Gurramkonda, Yanyan Qu, Dingyin Tao, Christopher A. LeClair, James Swartz, Govind Rao

## Abstract

Recent studies project that the prevalence of diabetes is expected to increase significantly and lead to escalating demand on the insulin supply chain. Despite being the first recombinant therapeutic approved by the FDA, insulin remains challenging to access for many around the globe. Here we report on advancements in manufacturing insulin using cell-free protein synthesis (CFPS) systems to rapidly produce mature desB30-insulin in less than a day. This is a major advance towards decentralizing insulin manufacturing and bringing production closer to the point-of-care, thereby improving diabetic patient accessibility. To this end, a purified cell-free extract, PUREfrex^®^ 2.1, was utilized to synthesize a tagged proinsulin construct that can be captured and converted into mature insulin using an on-column affinity chromatography process. Notably, two chaperones, peptidyl prolyl isomerase (FkpA), and seven kilodalton protein (Skp) were observed to play a critical role during cell-free expression of proinsulin. The proinsulin was then immobilized on a Ni-NTA column where the purification and conversion of cell-free products were performed sequentially to yield desB30-insulin. Following further optimization, this method serves as a time and resource-efficient production process as compared to current methods. When applied simultaneously, the cell-free expression and on-column conversion methods reported here can be adopted to enable rapid on-demand insulin manufacturing in order to improve the accessibility of insulin and prevent future shortages.

## Introduction

Globally, three major biopharmaceutical companies, Eli Lilly, Novo Nordisk, and Sanofi, control greater than 96% of the insulin market by volume [1,2]. This monopoly over insulin manufacturing has enabled a select few companies to dictate insulin prices in low-, middle-, and high-income countries [3,4]. As a result, a recent study showed that the mean accessibility of insulin in health facilities that should have insulin was as low as 55% for 13 low- and middle-income countries (LMIC) [5]. These barriers to access are not exclusive to LMIC, as evidenced by the inaccessibility of insulin in the United States, with a 2022 study reporting that 1.3 million diabetic Americans ration insulin due to cost [6]. To address this issue, several initiatives led by state and federal governments have sought to reduce the monthly cost of insulin [7,8]. While these initiatives assuage the individual burden of people living with diabetes, challenges around access are likely to increase as the global diabetic population is estimated to reach 643 million by the year 2030 [9]. Additionally, as evidenced by the COVID-19 pandemic, supply chain vulnerabilities can result in devastating consequences for the distribution of critical biologics, with insulin being no exception [10]. Therefore, the need for a rapid and decentralized insulin biomanufacturing approach is essential to ensure insulin access for individuals across the globe. Here, we discuss the expression of proinsulin using a purified cell-free protein synthesis (CFPS) system, PUREfrex^®^ 2.1, and the purification and “on-column” conversion of the precursor product to yield human desB30-insulin.

Recombinant human insulin has a variety of therapeutic analogs and is widely used for the treatment of Type I and Type II diabetes [11]. Structurally, insulin (5.8 kDa) is a 51 amino acid (AA) peptide hormone that contains two inter- and one intra-disulfide bond (A7-B7; A20-B19 and A6-A11, respectively), and is the post-translational product of proinsulin (~9 kDa, insulin with a C-peptide) [12]. The proper formation of these disulfide bonds occurs within the mammalian endoplasmic reticulum (ER) and is mediated by the C-peptide which is flanked by dibasic residues at each end (Arg-Arg and Lys-Arg) [13]. Upon transport to the Golgi, proinsulin is converted to native human insulin and C-peptide (~3 kDa) via protease digestion and stored in secretory vesicles. When increasing blood glucose levels signal the release of insulin, cellular glucose uptake is facilitated via glucose transporters like GLUT4. Additionally, insulin tends to form dimers and zinc-coordinated hexamers while stored in secretory vesicles [14].

Generally, utilization of large bioreactor-based facilities culturing recombinant organisms, typically of bacterial or yeast origin, has been the biotechnology standard since the first licensed insulin production process gained approval by Eli Lilly in 1982 [15,16]. In these first bioprocesses, proprietary cell lines were cultured to thousands of liters to express insulin A and B chains independently. Nowadays insulin is first produced as a precursor molecule in the form of proinsulin. These processes are generally referred to as the “two-chain” and “proinsulin” production approaches [17]. Proper disulfide bonding using the “two chain” approach resulted in low yields of properly folded insulin; thus the “proinsulin” approach is now utilized as it is more efficient [18,19]. For this reason, we selected the “proinsulin” approach as the basis for our bioprocess design. Typically, the purification and conversion of proinsulin into recombinant human insulin requires the solubilization of *E. coli* inclusion bodies (IB), which requires denaturation followed by oxidative sulfitolysis, proper formation of disulfide bonds, and enzymatic cleavage using one or more trypsin-like proteases [18]. The converted insulin is purified to greater homogeneity using a series of well-established chromatography steps including cation exchange (CEX), anion exchange (AEX), and reverse phase (RP) chromatography for removal of unwanted cleavage intermediates, DNA, endotoxin, and deamidated insulin. It is important to note that the challenges regarding insulin manufacturing are a result of the proprietary nature of these processes. Thus, the ability to provide a well-defined, open-source method for easy and reproducible insulin production could enable new manufacturers to enter the insulin market, effectively increasing competition and driving down selling prices to improve accessibility.

In this study, we develop an initial bioprocess that rapidly produces proinsulin using CFPS, followed by a combined purification and conversion step to produce recombinant human insulin. Given recent advances in distributed cell-free biomanufacturing, we believe this approach can be readily adapted with further development into an automated biomanufacturing platform, such as the Bio-MOD system [20,21]. Initially, several in-house and commercial cell-free extract-based systems were screened with little success, and this motivated investigation of PURE based CFPS. The PUREfrex^®^ 2.1 product was most successful at expressing proinsulin and is composed of 36 purified components from *E. coli* that are required for transcription and translation [22]. Additionally, the PUREfrex^®^ 2.1 system is void of proteases and peptidases which are particularly harmful to lower molecular weight proteins such as insulin. Given CFPS production is not bound to the confines of a cell wall, two periplasmic chaperones, FkpA, and Skp, were easily screened to evaluate their possible enhancement of proinsulin expression. These chaperones were previously reported to enhance soluble protein titer in the cell-free production of complex disulfide bonded proteins [23] and are periplasmic chaperones in *E. coli* [24–26]. Additionally, by utilizing these chaperones to express proinsulin in a soluble state, the need for proinsulin solubilization and refolding prior to purification is diminished [18]. This soluble expression then enables for effective capture and “on-column” conversion of cell-free expressed proinsulin, followed by enzymatic cleavage using trypsin or similar serine proteases to convert proinsulin into desB30-insulin, as outlined in Figure 1. To this end, we successfully expressed, converted, and purified recombinant proinsulin within a cell-free system while utilizing a novel approach to generate desB30-insulin.

**Figure 1.**
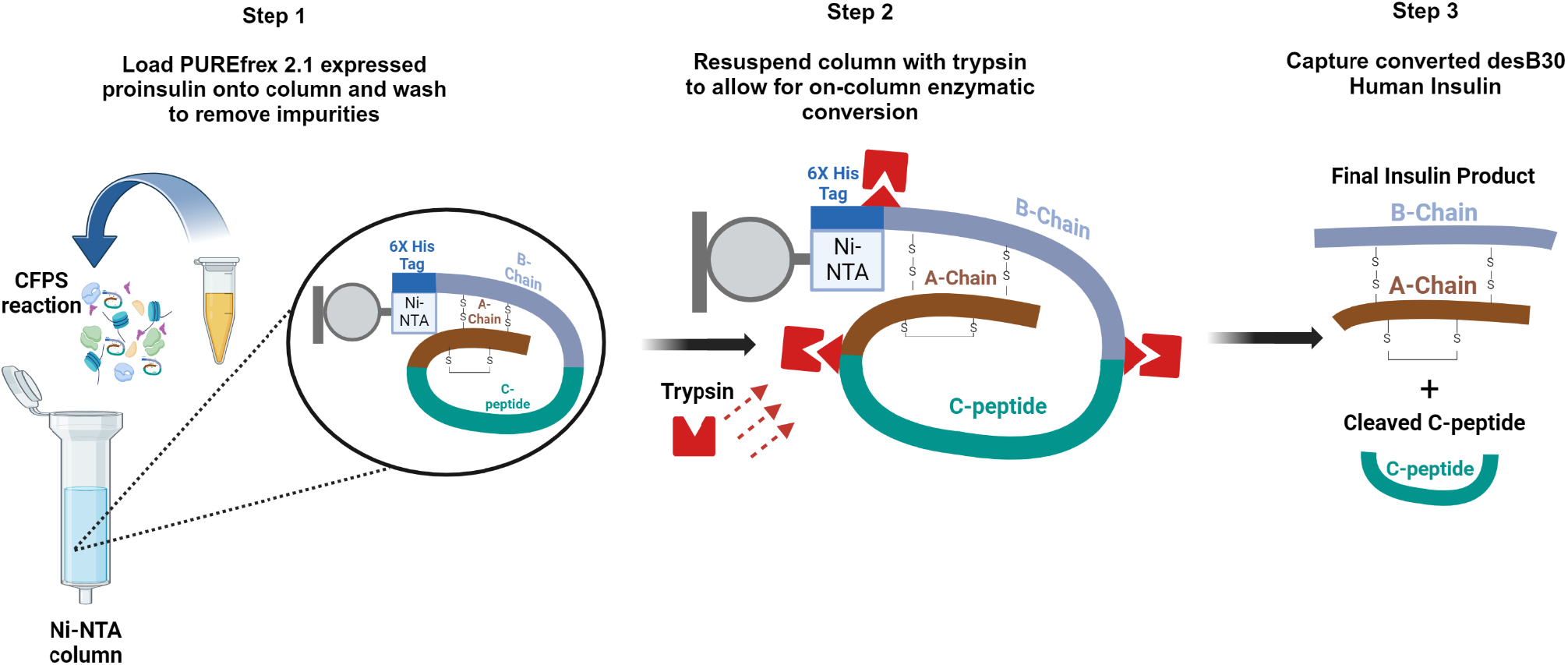
Proof of concept for on demand insulin production showcasing three major steps to produce and convert proinsulin into active insulin. Step 1 – proinsulin expression using PUREfrex^®^ 2.1 cell-free system with FkpA and Skp supplementation; Step 2 – capture and purify proinsulin using a Ni-NTA resin, then perform enzymatic conversion using trypsin to remove C-peptide and enable release of converted insulin from the column; Step 3 – collect insulin product and subject to additional downstream purification to meet purity requirements for therapeutic use.

## Material and Methods

### Proinsulin Gene Construction and DNA Preparation

The His-R-IP construct (10.85 kDa) was engineered from the native human proinsulin sequence (P01308) with minor alterations to amino acid composition (Supplementary Table 1). Since the desired gene of interest is intended to be produced via CFPS, emphasis was placed on minimizing mRNA secondary structures in the area of translational initiation. Codon variations to prevent secondary structure were evaluated using mFOLD software. The resulting genes were synthesized (Genscript, Piscataway, NJ, USA), then introduced via Gibson Assembly between a T7 promoter and T7 terminator in a pUC19-based vector (New England Biolabs (NEB), Ipswich, MA, USA), pMZL. DNA plasmid preparation for cell-free protein synthesis (CFPS) reactions was conducted using the Zymo Maxiprep Kit II (Cat# D4202) (Irvine, CA USA) using the recommended protocol with water as the elution buffer. Similar methodology to that described here was also used to develop a His-EK-IP construct (Supplementary Table 1), which contained an enterokinase cleavage site for the removal of a hexahistidine tag and was expressed in ALiCE^®^.

### PUREfrex^®^ 2.1 Expression

A commercially available purified cell-free extract, PUREfrex^®^ 2.1(GeneFrontier, Kashiwa, Japan), was used for the expression of His-R-IP and contains 36 purified components needed for transcription and translation [27]. The PUREfrex^®^ 2.1 system is preferred for assembling proteins with complex disulfide bonding given that it does not contain disulfide reduction activity found in other commercially available CFPS systems, thereby allowing the user to optimize precise concentrations of reduced (GSH) and oxidized (GSSG) glutathione based on the target protein. As previously reported, 8µM *E. coli* disulfide bond isomerase C (DsbC) in the presence of 4mM reduced (GSH) and 3mM oxidized (GSSG) glutathione were added for optimal disulfide bonding [28]. Peptidyl-prolyl isomerases (FkpA) and seventeen kilodalton periplasmic protein (Skp) were provided by Swartz Lab and supplemented into the reaction. Final concentrations of 25, 50, and 100 *μ*M FkpA, as well as 50, 100, and 200 *μ*M Skp were supplemented into the reaction. No chaperone and no water adjustment controls were also included to evaluate the benefit of FkpA and Skp and the effect of dilution of PUREfrex^®^ 2.1 reaction components. His-R-IP and His-EK-IP template DNA were prepared using a Zymo Maxiprep Kit II and added at a final concentration of 15 µg/mL. Expressions were performed at the 20 µL scale for 16 hr at 37 °C in 1.5 mL Eppendorf tubes in triplicate (n = 3) on a ThermoMixer C. Samples were then spun down at 10,000 g for 10 min and soluble products were analyzed via western blot analysis.

### SDS-PAGE Western Blot Analysis

Using 1.5 mL Eppendorf tube, 1-28 µl of 1X PBS was aliquoted and mixed with 2-29 µL of the sample. These diluted samples were then treated with 2X Reducing (~140 mM β-mercaptoethanol) Tris-Tricine (BioRad #1610739) or 6X Reducing (~210 mM β-mercaptoethanol) Laemmli (Boston Bioproducts, BP-111R) sample buffer, then boiled at 100 °C for 5min. A non-reducing version of each sample buffer was also used. Samples were spun at 16,000 x g for 1 min, loaded onto a 16.5% Tris-Tricine Gel or 4-20% Tris-Glycine gel, and run at 120 V for 2 hr or 200 V for 40 min, respectively. The gel was then cracked, and the gel was transferred into the blotting apparatus immersed in 1X Tris-glycine transfer buffer. A nitrocellulose membrane, had been submerged in transfer buffer for 5 min, was then placed on the gel apparatus, allowing for protein transfer onto the nitrocellulose membrane. Upon completion of protein transfer at 100 V for 45 min, the membrane was submerged in 20 mL of blocking solution (5% skim milk, 2% PVP in 1XPBST) for 1 hr. The blot was then probed with rabbit anti-insulin antibody (Cell Signaling, C27C9) at a 1:1000 dilution for 1 hr, followed by three washes (5 min each) of 1X PBST. The blot was then probed with goat anti-rabbit HRP-conjugated secondary antibody (Abcam, ab6721) at a dilution of 1:5000 for 1hr, washed with PBST, and then developed using SuperSignal West Dura (Thermo, 34075). Next, the membrane was imaged using a ChemiDoc XRS+ System and images were analyzed using ImageJ densitometry (BioRAD). A standard curve ranging from 250 to 1000 ng was generated using 100 µg/mL proinsulin (BioTechne, 1336-PN-050) and human insulin (USP, 1342106) standards for qualitative and quantitative analysis of expressed products. The resulting bands in the standard curve lanes were quantified using ImageJ densitometry and used to calculate the concentration in the unknown samples.

### SDS-PAGE Coomassie Staining

Gel loading samples were prepared and electrophoresed as per the Western blotting procedure. First, gels were washed three times with deionized water, 5 min each, then immersed in Coomassie Brilliant Blue R-250 solution (BioRad, #1610400) for 1 hr with gentle shaking. The gel was then de-stained using a buffer consisting of 50% deionized water, 40% methanol, and 10% acetic acid for a minimum of 2 hr until the background became clear. A standard curve ranging from 250– 1000 ng was generated using insulin (USP, 1342106) and proinsulin (Biotechne, 1336-PN) standard (100 µg/mL) for qualitative and quantitative analysis of conversion products. The resulting bands from the standard curve were quantified using ImageJ densitometry and used to calculate the concentration of the unknown samples.

### Purification of Proinsulin using Immobilized Metal Affinity Chromatography (IMAC)

For IMAC purification of components from the PUREfrex^®^ 2.1 CFPS system, purification of His-R-IP was achieved by using Ni-NTA resins (ThermoFisher, #88221). A calculated amount of resin was added to a 0.8 mL Pierce centrifuge spin column based on the estimated soluble protein yield obtained by Western blot analysis. Before protein binding, the resin was washed three times with five column volumes (CV) of 1X PBS, pH 7.4, and spun at 500 x g for 2 min. The harvest was then diluted with a binding buffer consisting of 1X PBS + 5 mM imidazole, pH 8.0 at a 1:4 (v/v) binding buffer to harvest ratio, then used to resuspend the Ni-NTA resin. Next, the sample was mixed on a rotary shaker for 20 min at room temperature. Samples were then spun down and flow-through (FT) was collected for future calculation of recovery. Post-loading impurities were washed by adding wash buffer (10 CV), 1X PBS + 35 mM imidazole, pH 7.4, and spinning at 500 x g for 2 min. The sample was then eluted four times with 5 CV elution buffer, containing 1X PBS + 300 mM imidazole, pH 7.4. Elution samples were analyzed using SDS-PAGE Coomassie Stains.

### “On-column” Enzymatic Conversion

For conversion, 15 µL of Ni-NTA was equilibrated with 1X PBS pH 7.4 for a total of three, 5 CV washes in a 0.8 mL spin column. The resin was then resuspended with a mixture containing 20 µg of PUREfrex^®^ 2.1 His-R-IP expressed with 100 µM Skp and then diluted with a binding buffer consisting of 1X PBS + 5 mM imidazole at pH 7.4 at a ratio of 1:4 (v/v). Next, the mixture was incubated with end-over-end mixing at room temperature for 20 min. The FT was collected, and the resin was washed with 10 CV of wash buffer (1X PBS + 35 mM imidazole, pH 7.4). A reconstituted trypsin solution (100 µg/mL) was prepared fresh by mixing 200 µL of 50 mM acetic acid with 20 µg of Promega Sequencing Grade Modified Trypsin and incubated at 30 °C for 15 min before addition “on-column”. A 1.5 µg/mL solution of trypsin containing 50 mM ammonium bicarbonate at pH 8.0, equivalent to 26 CV, was then prepared and mixed with the resin end-over-end at 37 °C for 2.5 hr. This resulted in a ratio of trypsin to proinsulin of 1:34 (w/w). The 2.5 hr “on-column” FT was collected and stored for further analysis. The column was then incubated with two 13 CV elutions (1X PBS + 300 mM imidazole, pH 7.4) for 2 min each to recover converted insulin following “on-column” conversion. Both 2.5 hr “on-column” FT and post “on-column” elutions were adjusted to pH 4.0 with acetic acid (<10 µL) to stop enzymatic activity. All conversion samples were subject to SDS-PAGE analysis. Percent recovery was calculated by multiplying the concentration of each sample by the total volume collected, then dividing by the total proinsulin harvest mass.

### High-Performance Liquid Chromatography-time-of-flight Mass Spectrometry (HPLC-TOF-MS)

High-performance liquid chromatography (HPLC) was performed on an Agilent 1290 Infinity II LC system (Agilent Technologies, Wilmington, DE, U.S.A.) equipped with a diode array detector (DAD), binary pump, multicolumn thermostat, and autosampler. The mobile phases used for the separation were MS-grade water with 0.1% formic acid (solvent A) and MS-grade acetonitrile with 0.1% formic acid (solvent B). Gradient elution of proteins from the analytical column (HALO BioClass Protein Diphenyl, 1000 Å, 2.7 µm, 2.1 x 100 mm) was performed using a gradient starting at 15% B at a flow rate of 0.3 mL/min. The mobile phase was then increased from 15 to 80% B over 8 min, and maintained at 80% B for 3 min, followed by a 80 to 15% decrease of B over 1 min and re-equilibration of the column with 15% B for 4 min. Separations were performed at a column temperature of 35 °C with a total run time of 16 min. Various volumes of sample were injected onto the column for each experimental run. Mass spectrometry (MS) experiments were conducted on an Agilent 6230 TOF system (Agilent Technologies, Wilmington, DE, U.S.A.), equipped with a DUAL AJS ESI source operating in positive ion mode. MS spectra were acquired from m/z 400 to 3200 at a scan rate of 1 spectrum per second with Profile format. The electrospray ionization (ESI) source parameters were used as follows: gas temperature, 300 °C; gas flow, 7 L/min; nebulizer, 35 psi; sheath gas temperature: 275 °C; Vcap, 4500 V; nozzle, 1000 V; fragmentor, 275 V. HPLC-TOF data analysis was performed by Agilent MassHunter Qualitative Analysis (B.07.00) with BioConfirm software (B.07.00) workflow installed for intact protein analysis. The parameters for protein deconvolution were set as: Deconvolution algorithm: Maximum Entropy, mass range: 4,000 to 50,000 with a mass step at 1.0 Da. Subtract baseline: baseline factor 7.0, adduct: Proton.

## Results

### PUREfrex® 2.1 Expression of Biosynthetic Proinsulin

Given that one of the central aims of this work is to perform post-translational processing of proinsulin while immobilized “on-column”, three protein elements, including the location of the affinity tag, design of a flexible linker peptide, and integration of site-specific enzyme recognition sequences within proinsulin were added, as outlined in Figure 2. First, the N-terminal signal peptide of pre-proinsulin was replaced with a hexahistidine tag (HHHHHH) since this region of precursor insulin is typically cleaved in vivo, and the introduction of hydrophilic amino acids has been shown to enhance the solubility of proinsulin [29]. Following the affinity tag, a glycine linker motif (GGGG) was added to enable flexibility of the affinity tag and improved capture during purification. For concurrent removal of the hexahistidine tag and C-peptide during “on column” conversion, an arginine linker (R13) was also introduced between the glycine linker motif and native proinsulin sequence. The design of this N-terminal extension provides the correct N-terminus for the final product [30]. Additionally, to ensure proper removal of the C-peptide during insulin production, the lysine at position 78 was replaced with alanine (K78A) and has been shown to enhance trypsin cleavage efficiency at position R79 by preventing the formation of the unwanted intermediate, arg-AO-insulin [31]. Insertion of an aspartic acid residue (D44) at the N-terminus of the C-peptide was also introduced to allow for Asp-N endoproteinase cleavage for generation of a native insulin B-chain upon enzymatic processing but was not utilized in the present study.

**Figure 2.**
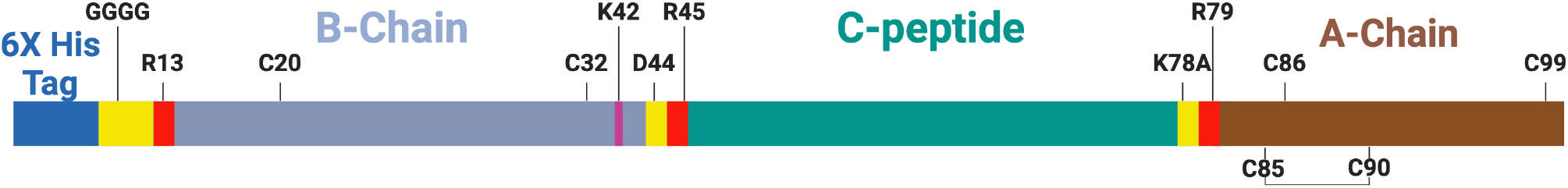
Design and functional features of the His-R-IP construct. Schematic of proinsulin construct used for “on-column” conversion into desB30-insulin. 6X His tag (dark blue) is fused via flexible glycine linker followed by an arginine at position 13 (R13) to enable for native insulin N-terminus. Native proinsulin containing B-chain (light blue), C-peptide (green), and A-chain (brown) were preserved except for an additional aspartic acid, D44, allowing for AspN endopeptidase cleavage. Additional mutation of lysine at position 74 to alanine, K78A, preventing premature cleavage at position 78 by Trypsin. Red denotes positions in which enzymatic cleavage is required for conversion into insulin. Yellow denotes positions which were altered/inserted into the native proinsulin sequence. Cysteines (C) which participate in disulfide bonding and their respective spatial placements within the construct are also provide.

Initial His-R-IP expression in KC6 extract (Supplementary Methods 1) resulted in no protein. This warranted sequence verification of the insert in the pMZL vector, for which 100% sequence fidelity to the designed insert was obtained. A subsequent expression was performed by pretreating the KC6 extract with 0.5 mM IAM for 30 min in the dark, followed by expression of pMZL His-R-IP (10, 15, and 20 ng/μL final DNA concentration) with a 1:4 ratio of reduced (GSH) to oxidized (GSSG) glutathione and 8 μM DsbC. Upon Western blotting of the harvest material, no protein was observed. Experimental conditions which failed to yield proinsulin expression are provided in Supplementary Figure 3. Furthermore, subsequent testing of these conditions was also performed in the NEBexpress (New England Biolabs) CFPS system which utilizes a similar cell-free extract and resulted in no protein expression (data not shown). Expression screening was continued using the PUREfrex® 2.1 system where 25, 50, and 100 μM FkpA, as well as 50, 100, and 200 μM Skp was supplemented. Figure 3a shows control reactions performed without chaperones and water additions and resulted in dimer concentrations of 26.5 and 81.6 μg/mL, respectively, with no monomer present in either sample. The results in Figure 3b show that 100 μM FkpA concentration resulted in a roughly 1.5 log reduction in dimer formation in comparison to 25 μM. The 50 μM condition of FkpA yielded the highest concentration of monomer at 20.4 μg/mL and dimer at 309.3 μg/mL. For Skp, roughly one log reduction in dimer concentration was observed when 200 μM Skp was supplemented in comparison to the 50 μM (Figure 3c). Notably, the highest concentration of monomer was observed when 100 μM Skp was supplemented into the reaction and yielded monomer and dimer concentrations of 35.1 μg/mL and 621.3 μg/mL, respectively. In Figure 3d, the combined addition of FkpA and Skp at 50 μM and 100 μM for each chaperone resulted in monomer concentrations of 23.8 μg/mL and 16.1 μg/mL, respectively. Additionally, dimer proinsulin concentrations for the 50 μM and 100 μM conditions resulted in 148.4 μg/mL and 17.7 μg/mL, respectively.

**Figure 3.**
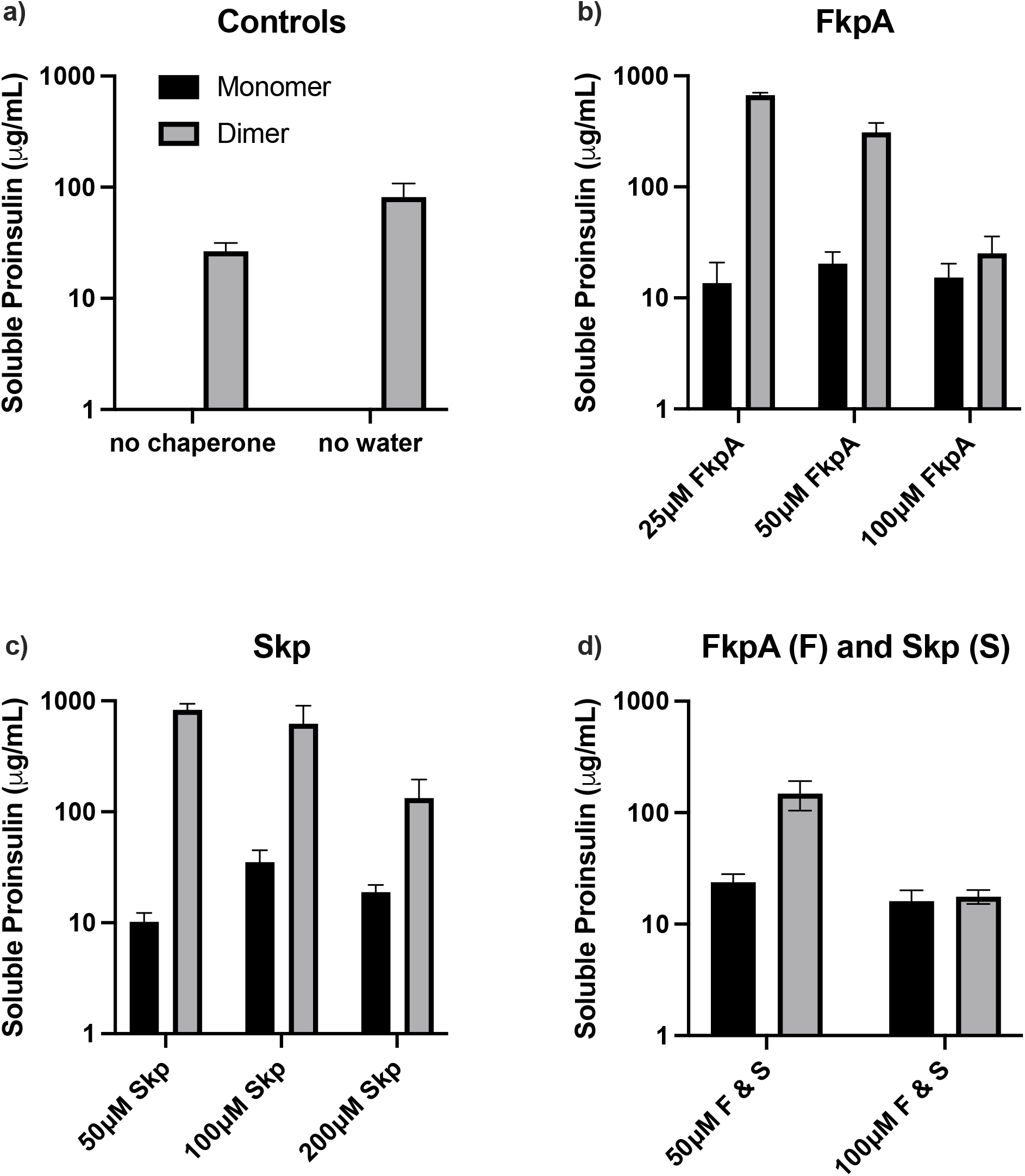
Supplementation of FkpA and Skp chaperones in PUREfrex® 2.1 for expression of pMZL His-R-IP and resulting soluble monomer (black) and dimer (grey) products obtained following expression. a) Positive controls with no chaperone and no water addition to the final reaction mixture; b) Base reaction with 25, 50, and 100 μM FkpA; c) Base reaction with 50, 100, and 200 μM Skp; d) A combination of FkpA and Skp supplemented at 50 and 100 µM, respectively. All reactions (20 uL) were incubated overnight (16 h) at 37 °C in 2 mL tubes in the presence of 8 μM DsbC, 4 mM GSH and 3 mM GSSG. Error bars represent the standard deviation (SD) of the mean (n = 3).

### Purification and Conversion of Proinsulin into Insulin

Using densitometry, roughly 56% of His-R-IP harvest was recovered following purification from PUREfrex® 2.1, yielding 11.2 μg of proinsulin immobilized “on-column” (Figure 4). Additionally, roughly 9.2% of His-R-IP was recovered in the form of desB30-insulin in the “on-column” products, with no proinsulin observed. Elution samples E1 and E2 in Figure 4 resulted in a total desB30-insulin mass of 1.1 μg, equivalent to an “on-column” process conversion recovery of 9.7%. Notably, a complex of proinsulin and insulin was observed at roughly 16 kDa and was verified using MS. This complex comprised roughly 41.9% of the elution product observed in both elution samples. The remaining percent of protein mass was in the form of un-cleaved proinsulin and intermediate side products.

**Figure 4.**
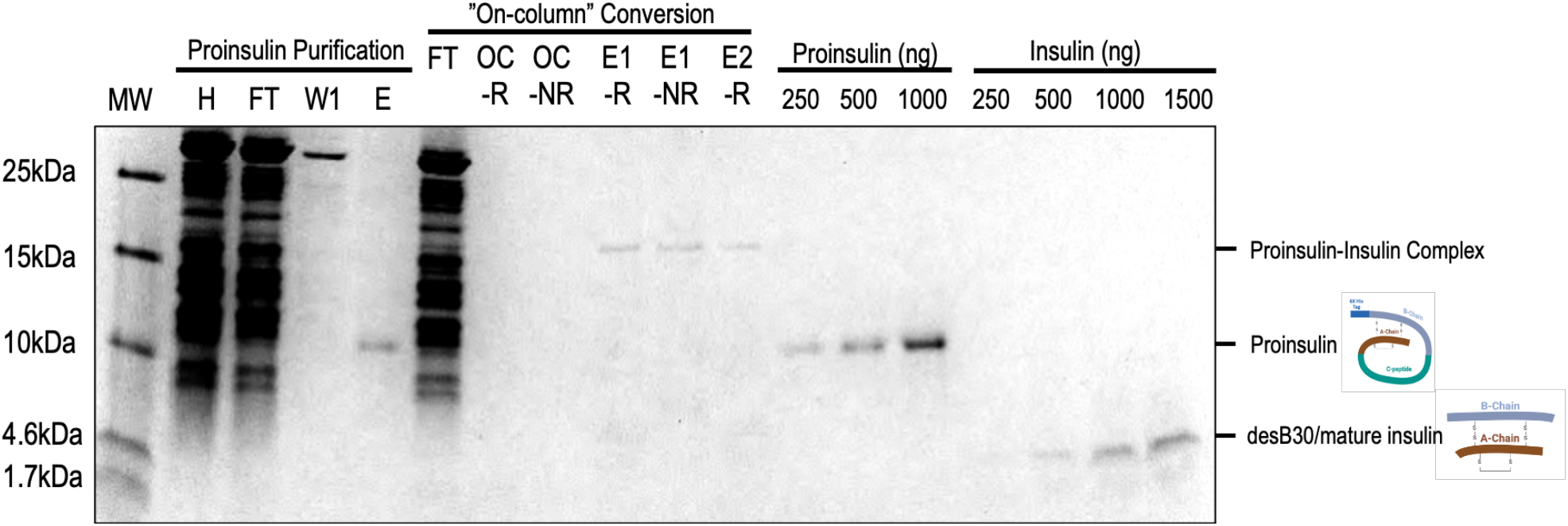
Coomassie stain of PUREfrex® 2.1 His-R-IP (proinsulin), “on-column” conversion products (proinsulin, insulin, C-peptide), proinsulin standard, and insulin standard. Harvest (H); flowthrough (FT); wash (W1); proinsulin elution (E); “on-column” FT products (OC); “on-column” elution fraction 1 (E1); “on-column” elution fraction 2 (E2); reducing conditions (R); non-reducing conditions (NR). Proinsulin – insulin complex (top notch, right) noted at approximately 16 kDa; Proinsulin noted at approximately 10.8 kDa; desB30-insulin noted at approximately 5.7 kDa. All elution and standard samples in this gel were subject to MS analysis (Figure 5) for determination of molecular identity and weight.

### Protein Characterization of Insulin and Proinsulin

The purified proinsulin and “on-column” elution samples from Figure 5 were subject to additional analysis using HPLC-TOF-MS for further characterization of protein identity and compared to molecular weight standards. The deconvoluted insulin and proinsulin standards (Figure 5a and 4b) were observed to be 5,807 Da and 10,471 Da, respectively, and were in commonality with their theoretical molecular weights of 5,807 Da and 10,476 Da. Figure 5c, which evaluated purified PUREfrex® 2.1 His-R-IP, showed two major peaks at 10,843 Da and 10,870 Da yielding a near match with the theoretical MW of the His-R-IP construct (10,848 Da). Semi-pure “on-column” elution fractions (E1 and E2) in Figure 5d resulted in a major peak at 5,717 Da, consistent with the theoretical molecular weight of desB30-insulin at 5,713 Da. Additionally, minor peaks at 10,869 Da and 16,774 Da were observed and consistent with the molecular weight of proinsulin and insulin/proinsulin heterodimers, respectively.

**Figure 5.**
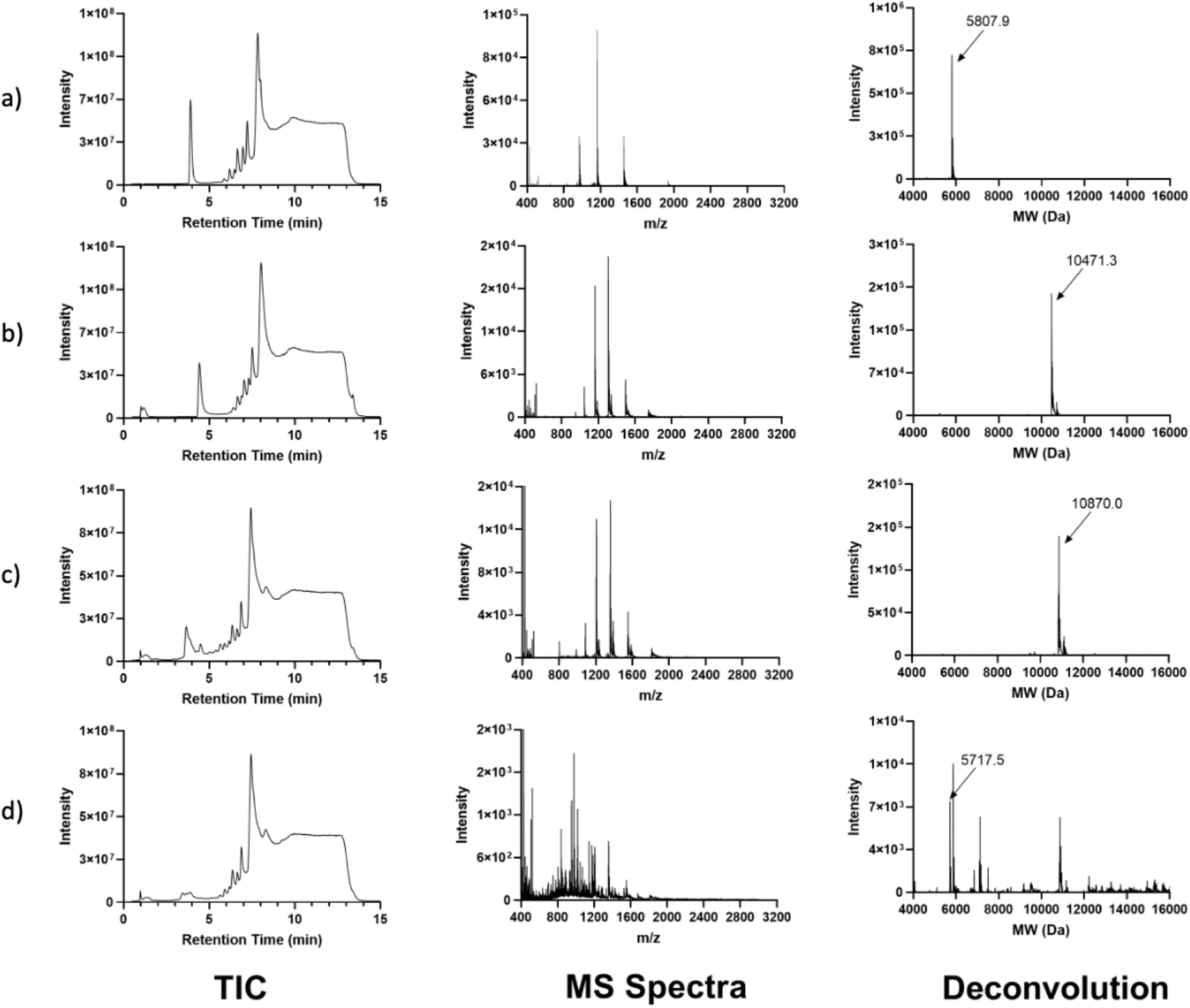
HPLC-TOF-MS Total Ion Chromatogram (TIC), MS spectrum, and deconvolution of (a) human insulin (MW 5,807 Da) and (b) proinsulin standards (MW10,471 Da) as well as (c) PUREfrex® 2.1 expressed His-R-IP proinsulin (MW 10,870 Da) following Ni-NTA purification. Additionally converted (d) desB30-insulin (MW 5,717 Da) from pooled “on-column” elution samples E1 and E2 from SDS-PAGE in Figure 4.

## Discussion

Our findings suggest that cell-free proinsulin expression is strongly dependent on specific chaperones typically found in the E. coli periplasmic space. In this study, the addition of 100 µM Skp to the PUREfrex^®^ 2.1 system resulted in soluble monomeric proinsulin expression at approximately 35.1 µg/mL, in contrast to the negative control, where no protein expression was observed (Figure 3). Additionally, supplementation of 50 µM FkpA resulted in a slightly lower yield of approximately 20.4 µg/mL. Interestingly, both chaperones tested resulted in substantially higher concentrations of dimer in comparison to the negative control, with yields of 621.3 µg/mL and 309.3 µg/mL, for Skp and FkpA respectively, following a reducing Western blot (Figure 3). Previous reports suggest that this dimerization of proinsulin is dependent on its intracellular concentration, with a dynamic equilibrium between monomers and dimers that interconvert in solution before folding occurs.[32]. Additionally, proinsulin’s resistance to low concentrations of reducing agents, such as DTT, has been previously reported, which may explain the failure of cell-free expressed proinsulin dimers to monomerize during reducing SDS-PAGE. (Figure 4) [32,33]

From this work, we propose that FkpA and Skp recruit exposed hydrophobic residues, such as those in regions A13-A19, A19-A20, B9-B19, and B20-23, which likely prevents aggregation of proinsulin. This interaction may slow folding kinetics, providing additional time for other chaperones and protein effectors to facilitate proper protein folding [34]. We speculate that during extract preparation, the concentration of periplasmic chaperones is greatly reduced, as the periplasm accounts for only 20-40% of a bacterial cell’s volume, and cell-free extracts are approximately 20-fold more dilute than intact cells [35,36]. Consequently, the concentrations of essential periplasmic chaperones, which are typically used during microbial production of proinsulin, are 95% lower in cell-free extracts compared to cell-based systems. The reduced availability of FkpA and Skp likely hinders the timely folding of proinsulin, promoting its misfolding. Misfolding is further exacerbated under higher proinsulin concentrations, leading to dimerization, as evidenced by the increased expression of proinsulin dimers in the no-water control (Figure 3a). This supports earlier reports that dimer formation is concentration-dependent and influenced by macromolecular crowding effects [32].

It is worth noting that proinsulin’s immediate access to periplasmic chaperones upon exiting the ribosome is a phenomenon possible only in a cell-free system, as in cells, access to these chaperones typically requires protein translocation to the periplasmic space. [37]. More importantly, co-chaperones such as the ones evaluated here are part of a more complex network of proteins that encounter nascent polypeptides as they emerge from the ribosomal tunnel [38–42]. As a result, proper modulation of their respective concentrations is critical for optimal protein folding and should be evaluated on a case-by-case basis. These concentration effects were observed in Figure 3d, where the combination of FkpA and Skp resulted in a reduction in both monomer and dimer proinsulin yields in comparison to each individual chaperone’s optimal concentration (Figure 3b and 3c). This suggested that adding higher concentrations of these holdases together could result in competition for hydrophobic regions within proinsulin and prohibit chaperones from staying attached long enough to enable proper folding.

The “on-column” purification and conversion results (Figure 4) suggest that expressed His-R-IP was converted to desB30-insulin within 2.5 hr, with faint bands observed at around 5.7 kDa for “on-column” elution fractions E1 and E2. While overall recovery was low at 9.7%, we believe optimization of trypsin to His-R-IP ratios (w/w) and improved buffering conditions, will enhance cleavage at the appropriate scissile bonds, thereby improving recovery [43,44]. As previously reported, the appearance of insulin in the “on-column” elution fraction suggests that desB30-insulin has a significant affinity for divalent cations such as Zn^2+^ and Ni^2+^ and requires imidazole for elution from the column [12,45]. In addition to the polyhistidine tag, enhanced coordination with Ni-NTA was likely motivated by HisB10 and GluB13 of the insulin metal binding site. As a result, this caused the insulin to remain bound despite cleavage of the N-terminal affinity tag. This contrasts with the intended bioprocess design, but it is encouraging given that the ability to coordinate divalent cations is an indication that the insulin is in the active T-state conformation [12]. This claim was further supported upon performing an insulin receptor binding assay where PUREfrex® 2.1 desB30-insulin displayed comparable binding activity to that of human insulin (Supplementary Figure 3). In future iterations of this work, adjustment of the N-terminal histidine tag to an alternative affinity tag should be made to allow for improved insulin release following “on-column” enzyme processing.

Analysis of samples was performed using HPLC-TOF-MS to identify the intact mass of the “on-column” elution products (Figure 5). The deconvoluted spectrum suggests that the “on-column” elution samples E1 and E2 (Figure 5D) yielded desB30-insulin with a MW of 5,717 Da, in close conformity to its theoretical MW of 5,713 Da. Notably, a 16,774 Da peak was also observed and is roughly equal to the sum of the MW for desB30-insulin and His-R-IP elution products (theoretical MW = 16,561 Da). Notably, the semi-pure elution of His-R-IP (theoretical MW = 10,848 Da) suggests that the N-terminal methionine is not removed during expression in the PUREfrex® 2.1 system, as evidenced by the 10,843 Da and 10,870 Da peaks in Figure 3C. This is likely due to the absence of methionine aminopeptidase (MAP) from the cell-free reaction mixture and proves not to be an issue for the “on-column” conversion process, as the N-terminal affinity tag is designed to be removed, yielding a native N-terminus [27,46].

While the results reported here show promise for producing insulin at the point-of-care, PUREfrex^®^ 2.1 and “on-column” conversion yields require further optimization to achieve the therapeutic daily dose of insulin, equivalent to 40 units/day (or 1.38 mg/day) [1]. To see if other CFPS systems could enable higher yields of monomer proinsulin expression, we also expressed His-R-IP in the ALiCE^®^ (*N. tabacum*) cell-free extract, which was capable of producing more than 320 µg/mL of soluble proinsulin in 16 hr at the 50 µL scale (Supplementary Figure 1A). Notably, it was found that shorter incubation times were beneficial to soluble product accumulation. This is likely due to the susceptibility of proinsulin to degradation, hence digestion by proteases is limited for incubations that are shorter and subsequently have less time for proteolytic activity (Supplementary Figure 1C) [47,48].

It was found that soluble protein accumulation was abolished at the 250 µL scale, suggesting that surface area to volume ratios play a significant role in protein synthesis as previously described (Supplementary Figure 1C) [49–51]. In addition to the proteolytic degradation discussed above, suboptimal oxygenation of lysate results in lower protein titers, resulting in no net protein yield. It should be noted that by leveraging ATP regeneration machinery such as the mitochondria in ALiCE^®^ (or inner membrane vesicles (IMVs) produced during *E. coli* lysate preparation), it is ensured that ATP is not rate limiting for optimal chaperone activity [51]. This may be one reason why the observed yields in ALiCE^®^ were higher than those observed in the PUREfrex^®^ 2.1 system, whose cell-free system does not contain any energy regeneration pathways (Figure 3). Therefore, ALiCE^®^ demonstrates the potential to produce milligram quantities of His-R-IP which can then be subject to conversion into insulin. Ultimately, the yields observed using ALiCE^®^ support the production of insulin in a portable biomanufacturing platform such as the Bio-MOD system, which has been configured to include cation exchange polishing procedures commonly used in the downstream purification of insulin products [52–56].

As advances in lyophilization and stabilization of cell-free extracts become more mainstream [57,58], we believe that the bioprocess technology outlined in this report will support future demand for insulin across diverse patient populations. Namely, the ability to express, convert, and purify insulin from a lyophilized cell-free system will enable on-demand production for patients in need and improve sustainability by reducing the amount of unused insulin. Additionally, this bioprocess approach could be adapted to produce a variety of different insulin analogs and enable rapid screening of insulin products with enhanced pharmacokinetics, such as smart insulins [59]. Thus, the expression, purification, and conversion procedures outlined in this report provide a promising alternative to current time-intensive production approaches and could pave the way for future decentralized insulin production.

## Acknowledgements

This work was inspired by two DARPA funded programs, Biologically Derived Medicines on Demand and Reimagining Protein Manufacturing. Encouragement and early discussions with Dr. Leah Tolosa-Croucher were helpful. Dr. Gurusingham Sittampalam for helping to establish valuable collaborations between UMBC and NCAT, without which our analytics would not have been possible; the late Dr. Antonio Moreira for emphasizing the importance of understanding the critical quality attributes of insulin during bioprocess design; Dr. Douglas Frey for purification guidance; and LenioBio for providing the ALiCE Lysate.

