## Supplementary Material for "On-demand insulin manufacturing using cell-free systems with an “on-column” conversion approach"

### Supplementary Documents

**Supplementary Methods 1. KC6 Extract Based Proinsulin Expression.** A KC6 *E. coli* extract was prepared as previously reported [60] and pre-treated with 500 µM iodoacetamide (IAM) for 30 min at room temperature on a rotator before the start of the cell-free reaction. Reduced (GSH) and oxidized (GSSG) glutathione were added to the reaction at concentrations ranging from 0-4 mM. Additionally, all reactions contained DsbC at a concentration of 8 µM. Chaperones Fkpa, Skp, PDI, and EroI, were also added at micromolar concentrations to evaluate their constituent effects on protein yield and solubility and were provided by the Swartz Lab. Cysteine was added into the reaction at 0.5, 1, and 2 mM to ensure the availability of cysteine was not rate-limiting during synthesis. All reaction formats assembled are outlined in Supplementary Figure 2.

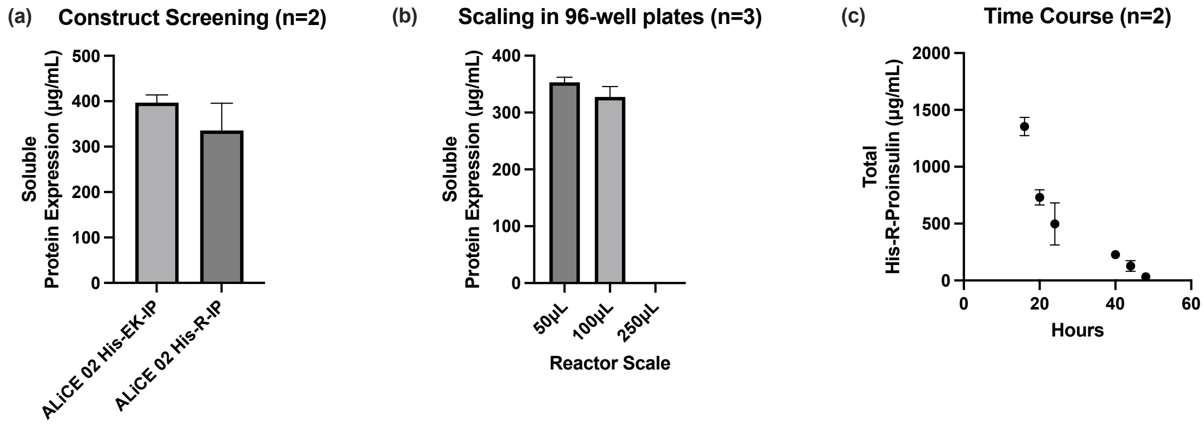


**Supplementary Figure 1.** (a) Initial Screening of Proinsulin constructs in ALiCE® – Western Blot Densitometry. Expression of all constructs in S.1 to establish a baseline. Reactions were performed at the 50 µL scale in 96-well plates for overnight expression at 25 °C with 750 RPM. Error bars represent the SD of the mean (n = 2). (b) Effect of Expression Scale on Soluble His-R-IP Titers – Western Blot Densitometry. Reactions were performed at the 50, 100, and 250 µL scale in 96-well plates for overnight expression at 25 °C with 750 RPM. Error bars represent the SD the mean (n = 3). (c) Time-course study on His-R-IP Construct – Western Blot Densitometry. Reactions were performed at the 50 µL scale in 2 mL tubes at 25 °C with 500 RPM for 16, 20, 24, 40, 44, and 48 hr. Error bars represent the SD of the mean (n = 2).

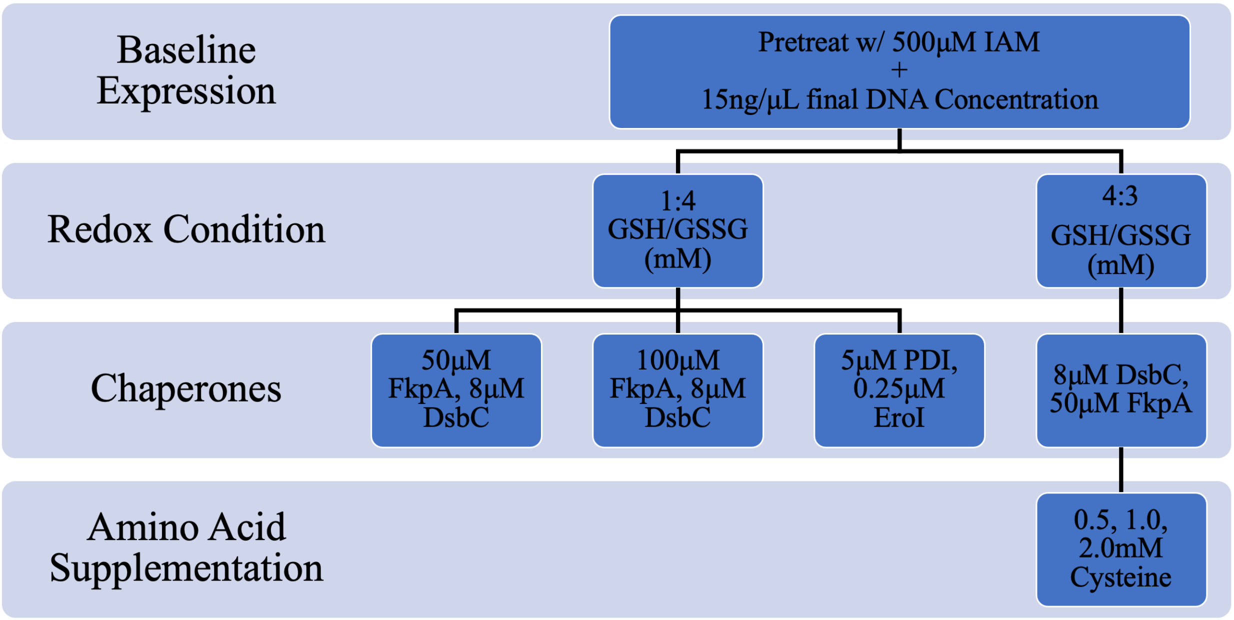


**Supplementary Figure 2.** Reaction conditions tested using KC6 *E. coli* CFPS system for expression of His-R-IP construct. All conditions contained IAM pretreated KC6 extract and then were subjected to adjusted redox environments, protein effectors, and amino acid conditions outlined - branching lineage indicates carryover of the prior condition. No protein expression was observed after Western blotting for any of the conditions outlined.

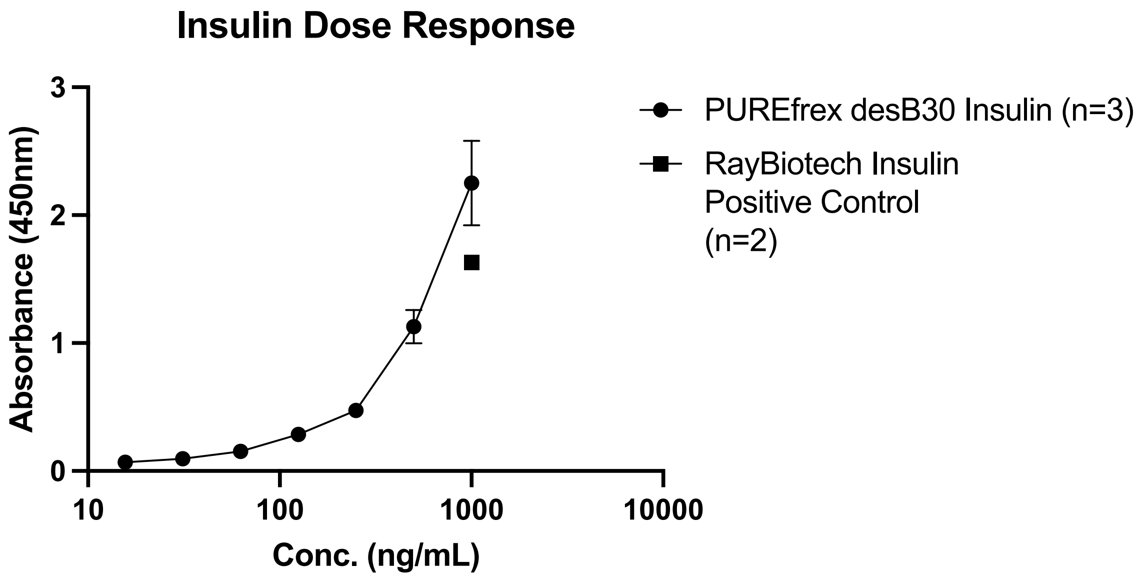


**Supplementary Figure 3.** RayBiotech Human Insulin / Insulin Receptor Binding Assay Kit (Cat# BAH-INSR-INS-1) was performed without receptor inhibition for determination of insulin binding affinity to insulin receptor. A titration of PUREfrex desB30-insulin post "on-column" conversion from 0 to 1000 ng/mL (n = 3) was performed and compared to RayBiotech insulin positive control at 0 and 1000 ng/mL (n = 2). Error bars display standard deviation (SD) from the mean.

**Supplementary Table 1.** PUREfrex® 2.1 Proinsulin Expression.

| Construct | MW (kDa) | Proinsulin Amino Acid Sequence |
|---|---|---|
| YSS-His-SS-EK-IP "His-EK-IP" | 11.678 | MYSSHHHHHHSSDDDDKFVNQHLCGSHLVEALYLVCGERGFFYTPKTDDRREAEDLQVGQVELGGGPGAGSLQPLALEGSLQKRGIVEQCCTSICSLYQLENYCN |
| G-His-GGGGR-IP "His-R-IP" | 10.848 | MGHHHHHHGGGGRFVNQHLCGSHLVEALYLVCGERGFFYTPKTDRREAEDLQVGQVELGGGPGAGSLQPLALEGSLQARGIVEQCCTSICSLYQLENYCN |
